# The unexplored diversity of wild lupins provides rich genomic resources and insights into lupin evolution

**DOI:** 10.1101/2024.03.07.583883

**Authors:** Karolina Susek, Edoardo Franco, Magdalena Tomaszewska, Magdalena Kroc, Humaira Jamil, Umesh Tanwar, Matthew N. Nelson, Roberto Papa, Massimo Delledonne, Scott A. Jackson

## Abstract

Lupin crops provide nutritious seeds as an excellent source of dietary protein. However, extensive genomic resources are needed for the adaptation of lupin crops, particularly to improve their nutritional value and facilitate their adaptation to harsh environments caused by the changing climate. Such resources can be derived from crop wild relatives, which represent a large untapped source of genetic variation for crop improvement. Here we describe the first whole-genome sequences of the cross-compatible species *Lupinus cosentinii* (Mediterranean) and its pan-Saharan wild relative *L. digitatus*, which are well adapted to drought-prone environments and partially domesticated. We found that both species are tetraploids, with similar genome structures, distributions of gene duplications, and numbers of expanded and contracted gene families. The expansion and contraction of gene families that determine seed size, a paradigmatic domestication trait, indicates that gene duplication may have led to morphological adaptations in *L. cosentinii* and *L. digitatus* differing from those in *L. albus*, a domesticated lupin used as a reference. Seed size may therefore reflect convergent evolution mechanisms that play a key role in lupin domestication.

## Main

The genus *Lupinus* (lupins) is part of the genistoid clade of leguminous plants (Fabaceae). It includes five domesticated but underutilized crop species that produce highly nutritious seeds (Nartea et al. 2023) with a protein content of up to 40% (Rawal 2019, Zhao et al. 2022). Lupins could therefore contribute to a healthy and sustainable human diet (Bellucci et al. 2021, Bulut et al. 2023, FAO 2023). The development of genomic tools can facilitate breeding and pre-breeding processes by exploiting the rich diversity of domesticated lupin species and wild relatives (Bohra et al. 2022). However, the number of whole-genome sequences available for the wild relatives of legume crops is limited to peanut (*Arachis hypogaea*), *A. duranensis* and *A. ipaensis* (Bertioli et al. 2016), soybean (*Glycine max*) and *G. soja* (Stupar 2010), and mungbean (*Vigna radiata* var*. radiata*), *V. reflexo-pilosa* var. *glabra* and *V. radiata* var. *sublobata* (Kang et al. 2014).

Whole-genome sequences have been published for two lupin crops, namely *L. angustifolius* (Hane et al. 2017) and *L. albus* (Hufnagel et al. 2020)(Hufnagel et al. 2020) providing some insight into lupin genome structure, diversity and evolution, but information from wild relatives is needed to take full advantage of lupin genetic resources. The genetic diversity of lupins has been highlighted by chromosome number and genome size (Naganowska et al. 2003, Susek et al. 2016) as well as epigenomic (Susek et al. 2017) and phylogenetic analysis (Aïnouche and Bayer 2004), and more recently the development of pan-genomes for *L. albus* (Hufnagel et al. 2021) and *L. angustifolius* (Garg et al. 2022).

There are ∼275 lupin species conventionally divided into ‘New World’ and ‘Old World’ types. Most annual and perennial lupins belong to the New World group and are found mainly in North America and the Andes, but only one species has been domesticated (*L. mutabilis*). The Old World lupins comprise ∼15 species distributed around the Mediterranean basin as well as North and East Africa (Gladstones 1998), but four of them are domesticated: smooth-seeded *L. albus*, *L. angustifolius* and *L. luteus*, and rough-seeded *L. cosentinii*. The somatic chromosome number of Old Word lupins varies widely (2*n* = 32, 36, 38, 40, 42, 50 or 52), with a basic chromosome number of *x* = 5–13. In contrast, most New World lupins have the somatic chromosome number (2*n* = 36 or 48), the exceptions being *L. bracteolaris*, *L. linearis* (2*n* = 32), *L. cumulicola* and *L. villosus* (2*n* = 52), but the basic chromosome number is proposed to be *x* = 6 in all cases. Multiple chromosome rearrangements have occurred among the Old World lupins, revealing a complex evolutionary process that suggests polyploidy (Susek et al. 2019). Indeed, there is evidence that *L. albus* and *L. angustifolius* evolved by genome duplication and/or triplication (Hane et al. 2017, Hufnagel et al. 2020, Kroc et al. 2014, Susek et al. 2019) from a diploid ancestor (Xu et al. 2020). These evolutionary processes not only involved chromosome rearrangements (Susek et al. 2019, Susek et al. 2016) but also epigenetic changes (Susek et al. 2017), and processes such as dysploidy and aneuploidy (Steele et al. 2010, Doyle 2012), suggesting that allopolyploidization may be unique to the Old World lupins (Drummond 2008). The genistoid clade, which diverged from the papilionoids soon after the emergence of legumes about 56 Ma (Gepts et al. 2005, Lavin et al. 2005), shows the highest frequency of polyploidy but is poorly characterized and only weakly supported as a sister clade to the remaining core papilionoids (Doyle 2012). Early data indicated that the genistoid basic chromosome number was *x* = 9 and the most common somatic chromosome number was 2*n* = 18 (Cannon et al. 2015, Goldblatt 1981).

Here, we investigated the genomic structure of two Old World lupin species: *L. cosentinii* Guss. (2*n* = 32) and *L. digitatus* Forsk. (2*n* = 36). Both are recognized for their drought tolerance, and *L. digitatus* serves as a source of drought-tolerance genes (Gladstones 1970). *L. cosentinii* Guss. is native to the western Mediterranean coast but has been introduced in Austria, Romania, South Africa and several parts of Australia (Gladstones 1998, Kole 2011). Furthermore, *L. cosentinii* cultivars Gus and Erregulla were domesticated in Australia from local wild germplasm in the twentieth century, and have improved traits such as low-alkaloid seed, non-shattering pods, early flowering, and soft seeds (Kole 2011). *L. cosentinii* was also crossed with *L. digitatus* (Gladstones 1998, Gupta et al. 1996), which is native to the pan-Saharan region (Gladstones 1998), and with another rough-seeded species, *L. atlanticus* (Gupta et al. 1996, Roy and Gladstones 1985). Interspecies crosses among the rough-seeded lupins revealed that the most compatible species are *L. cosentinii*, *L. digitatus* and *L. atlanticus*. Interestingly, *L. digitatus* seeds have been found in tombs of Egyptian Pharaohs, suggesting that domestication began more than 4000 years ago (Gresta et al. 2017). Seed size is a major domestication target in legumes because it is a key yield component and larger seeds can produce larger and more competitive seedlings. Also, evolutionary success of flowering plants depends on the seeds, thus seeds could be potentially used as a model to insight into evolutionary events, across different environments and ecological contexts. Our work provides a set of tools to facilitate the development of improved, climate-friendly lupin crops by exploiting the genetic diversity of wild species to promote the conservation and sustainable utilization of lupins as a source of dietary protein, and to increase the possibility of domesticating a greater variety of wild species.

## Results and discussion

### First wild lupin species whole-genome sequences: *de novo* genome assembly

We used a combination of techniques to generate the first genome assemblies for the wild, rough-seeded lupin species *Lupinus cosentinii* (*2n* = 32) and *Lupinus digitatus* (*2n* = 36). First, we produced PacBio HiFi reads (**Supplementary Table S1, Figure 1**) with ∼55× coverage for *L. cosentinii* and ∼43× coverage for *L. digitatus*. We then used *HiCanu* to generate assemblies of 650 and 492 Mb for *L. cosentinii* and *L. digitatus*, respectively (**Supplementary Table S2**). The assemblies were polished using 39 and 37 Gb of Illumina 150PE reads (**Supplementary Table S1**). This purging procedure reduced the assembly size to 588 Mb in *L. cosentinii* and 435 Mb in *L. digitatus* (**Supplementary Table S2**). We applied two sequential approaches to scaffold the contigs, first with 560 Gb (*L. cosentinii*) and 722 Gb (*L. digitatus*) of Bionano optical maps, then with 60.4 Gb (*L. cosentinii*) and 53 Gb (*L. digitatus*) of chromosome-level Illumina Hi-C data (**Supplementary Table S1**). The resulting *L. cosentinii* genome (588 Mb) was composed of 19 scaffolds (∼426 Mb, ∼72% of the assembled genome) and 709 further contigs, whereas the *L. digitatus* genome (435 Mb) consisted of 22 scaffolds (∼378 Mb, ∼87% of the assembled genome) and 339 remaining contigs (**Table 1**, **Figure 2a**). Benchmarking universal single-copy orthologs (BUSCO) reported 96.3% and 95.3% completeness with 23.6% and 22.5% duplicated genes in *L. cosentinii* and *L. digitatus*, respectively. The *L. cosentinii* genome was therefore larger than that of *L. albus* (451 Mb), which was similar in size to the *L. digitatus* genome. Depending on the species, lupin genomes are therefore similar in size to those of the common bean *Phaseolus vulgaris* (∼580 Mb) and the model legumes *Medicago truncatula* and *Lotus japonicus* (both ∼ 470 Mb).

**Figure 1:**
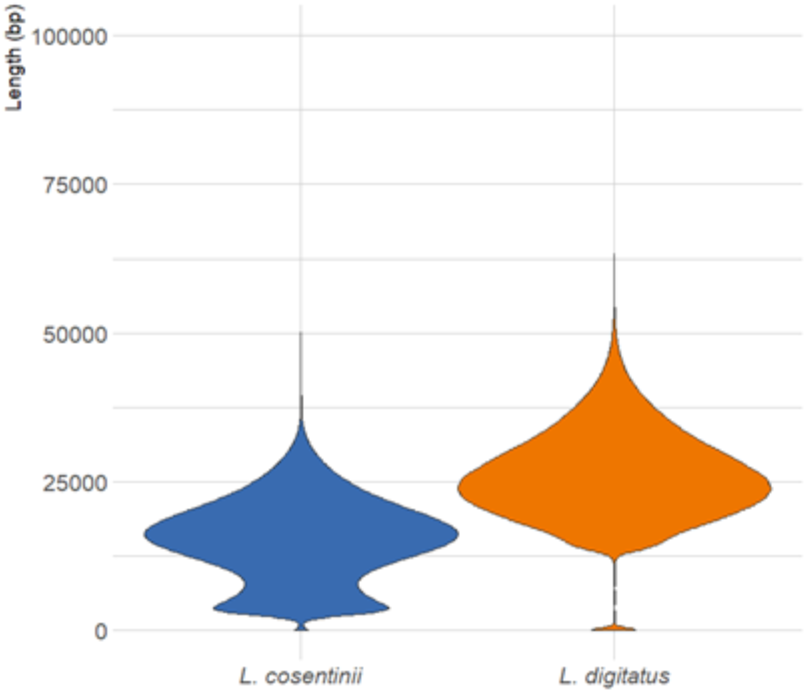
Length distribution of PacBio sequencing reads for *Lupinus cosentinii* and *L. digitatus*.

**Figure 2:**
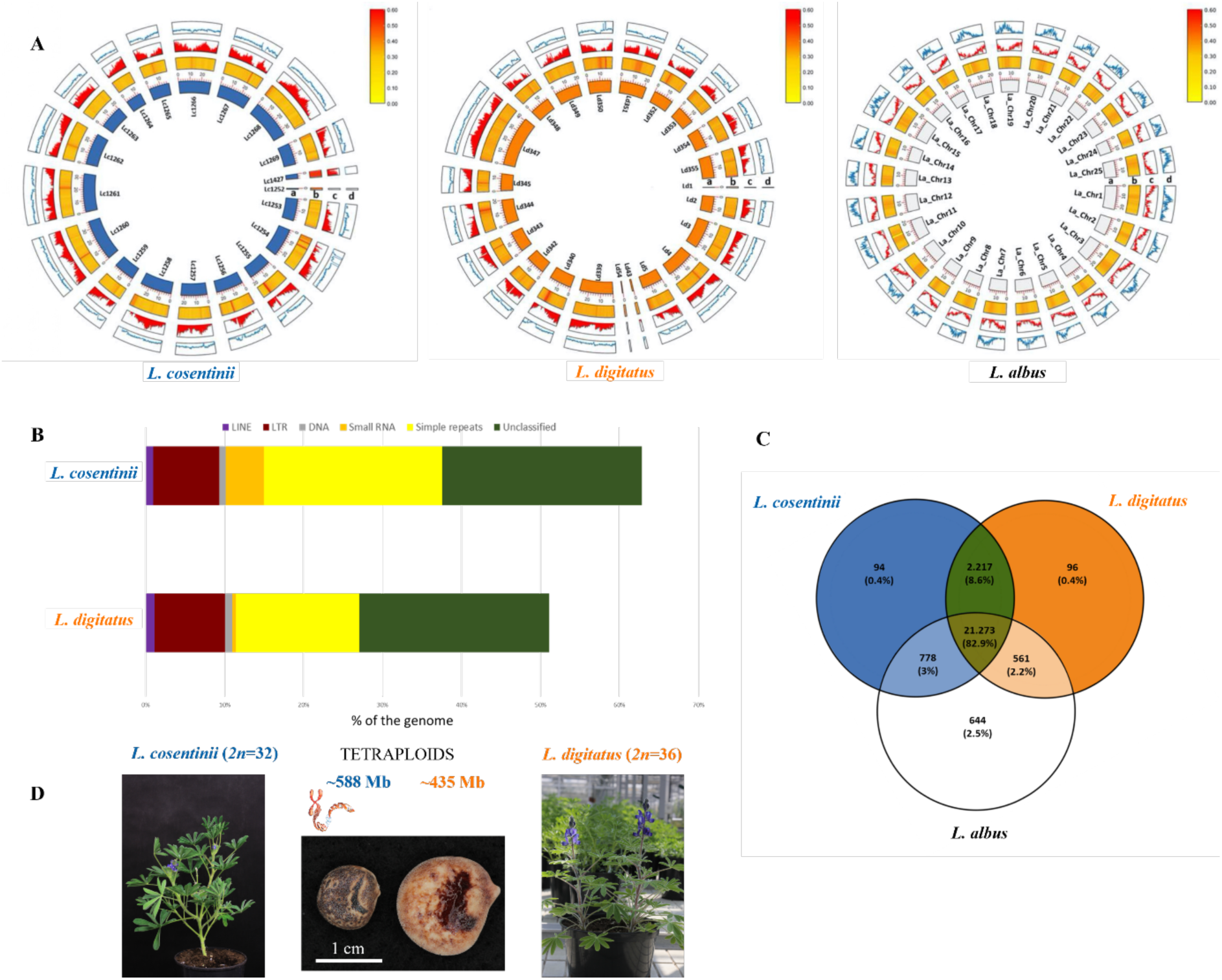
Genome assemblies and annotations of *Lupinus cosentinii*, *L. digitatus* and *L. albus*. The inner bars represent (a) the scaffolds (19 in *L. cosentinii* and 22 in *L. digitatus*) and the 25 chromosomes in *L. albus*. (b) GC content along the genomes. (c) Gene density. (d) Repeat element density. For ease of comparison, only scaffolds from *L. cosentinii* and *L. digitatus* were represented here (A); Sequence characteristics of repetitive DNA content in *L. cosentinii* and *L. digitatus* genomes (B); Venn diagram showing the number of gene families shared between *L. cosentinii* and *L. digitatus*, with *L. albus* as a reference (C); *L. cosentinii* and *L. digitatus* plants with their seeds (D).

**Table 1:**
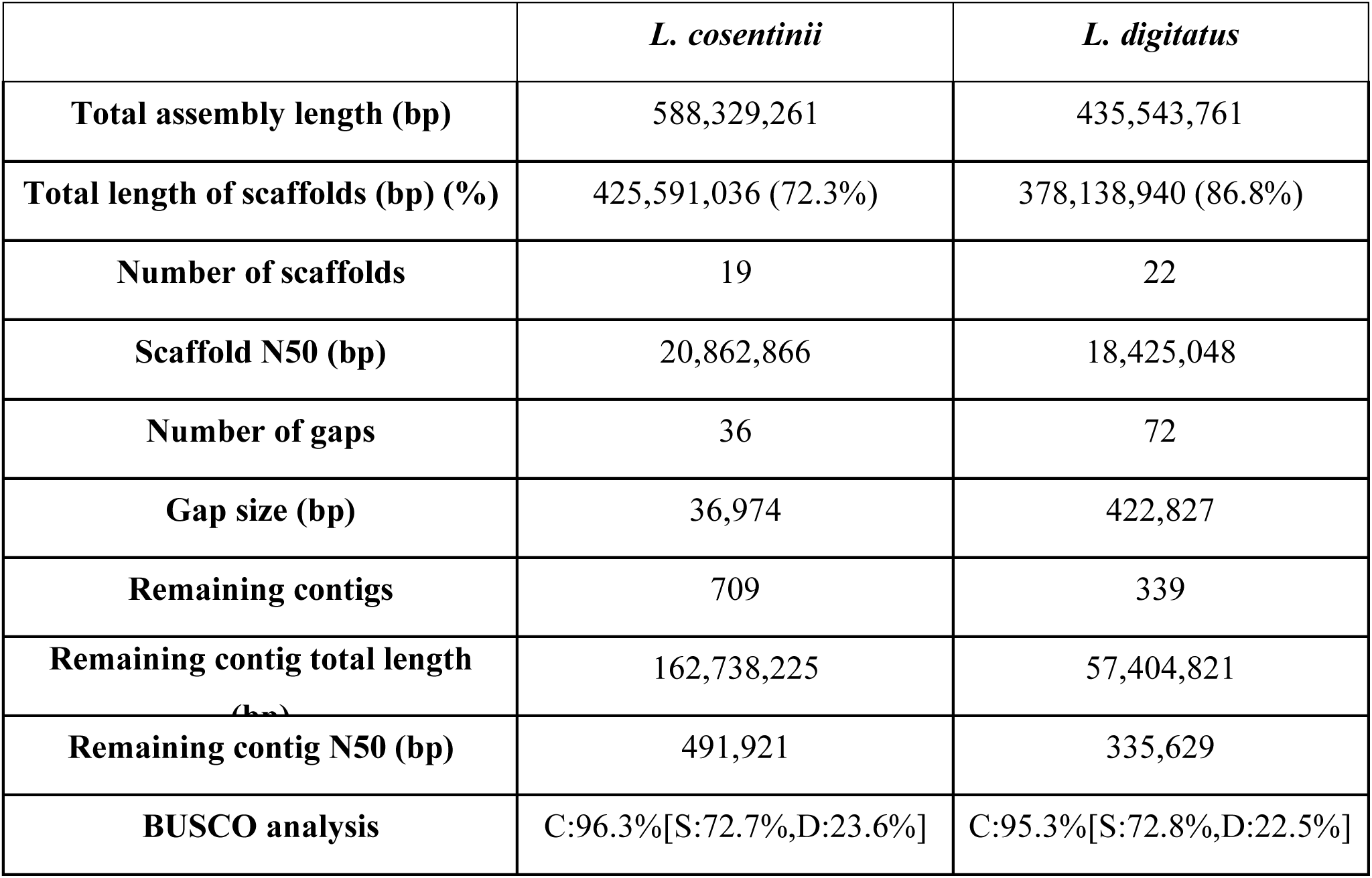
Summary statistics of the final *Lupinus cosentinii* and *L. digitatus* genome assemblies.

|  | *L. cosentinii* | *L. digitatus* |
| --- | --- | --- |
| **Total assembly length (bp)** | 588,329,261 | 435,543,761 |
| **Total length of scaffolds (bp) (%)** | 425,591,036 (72.3%) | 378,138,940 (86.8%) |
| **Number of scaffolds** | 19 | 22 |
| **Scaffold N50 (bp)** | 20,862,866 | 18,425,048 |
| **Number of gaps** | 36 | 72 |
| **Gap size (bp)** | 36,974 | 422,827 |
| **Remaining contigs** | 709 | 339 |
| **Remaining contig total length (bp)** | 162,738,225 | 57,404,821 |
| **Remaining contig N50 (bp)** | 491,921 | 335,629 |
| **BUSCO analysis** | C:96.3%[S:72.7%,D:23.6%] | C:95.3%[S:72.8%,D:22.5%] |

### Gene structure and composition of repetitive sequences

We predicted the presence of 34,780 and 31,260 genes in the *L. cosentinii* and *L. digitatus* genomes, respectively. For 26,860 (77.2%) and 25,478 (81.5%) of these genes, functional annotations were also present in high-confidence databases (SwissProt, RefSeq and TAIR) (**Supplementary Tables S3, S4**) and the *L. albus* proteome (**Table 2a**). Functional annotation with Gene Ontology (GO) terms was possible for 23,544 *L. cosentinii* genes (67.7%) and 23,019 *L. digitatus* genes (73.6%). Repetitive DNA accounted for 370.6 Mb (63%) and 224.3 Mb (51.5%) of the *L. cosentinii* and *L. digitatus* genomes, respectively. The major classes of repetitive elements were simple repeats, representing 22.5% and 15.5% of the *L. cosentinii* and *L. digitatus* genomes, and long terminal repeat retroelements (LTR-REs), representing 8.33% and 8.85%, respectively (**Table 2b**, **Figure 2b**, **Supplementary Table S5)**. For comparison, *L. albus* has a similar repeat content (60%) and also a low abundance (0.8%) of DNA transposons (Hufnagel et al. 2020).

**Table 2:** Characteristics of the *Lupinus digitatus* and *L. cosentinii* genomes. (a) Annotated genes. (b) Repetitive DNA.

| (a) | *L. cosentinii* | *L. digitatus* |
| --- | --- | --- |
| **No. of predicted genes** | 34,780 | 31,260 |
| **BUSCO completeness** | C:94.4%[S:71.4%,D:23.0%] | C:93.4%[S:71.1%,D:22.3%] |
| **No. proteins with functional annotation (% of total predicted)** | 26,860 (77.2%) | 25,478 (81.5%) |
| **No. proteins with GO description (% of total predicted)** | 23,544 (67.7%) | 23,019 (73.6%) |
| **No. proteins with GO description or a functional annotation (% of total predicted)** | 30,729 (88.4%) | 28,298 (91.8%) |
| (b) | *L. cosentinii* | *L. digitatus* |

|  | Length (bp) | Ratio (%) | Length (bp) | Ratio (%) |
| --- | --- | --- | --- | --- |
| LINE | 5,669,452 | 0.96 | 4,907,250 | 1.13 |
| LTR | 49,005,638 | 8.33 | 38,558,382 | 8.85 |
| DNA | 4,802,283 | 0.82 | 4,012,265 | 0.92 |
| Small RNA | 28,235,108 | 4.8 | 2,024,446 | 0.46 |
| Simple repeats | 132,281,679 | 22.48 | 67,524,865 | 15.5 |
| Unclassified | 148,443,427 | 25.23 | 104,045,282 | 23.89 |
| Repetitive content in total | 370,597,827 | 62.98 | 224,255,390 | 51.48 |

### Consequences of polyploidy during lupin evolution

We anticipated that *L. cosentinii* and *L. digitatus* would show some degree of polyploidy, like other legumes (Zhu et al. 2005). Accordingly, Genomescope/Smudgeplot indicated that both species are tetraploid (**Figure 3b-d**). A high degree of homozygosity (∼99.94%) was evident for both species, as shown by the single major peak in the *k*-mer distribution (**Figure 3a-c**; Kajitani et al. 2014). The distribution of biallelic single nucleotide polymorphisms (SNPs) in the genome assemblies of both species also indicated tetraploidy. The delta log-likelihood scores, calculated from the difference between the free model and the diploid, triploid and tetraploid models, were 1,202,737, 896,833 and 316,770, respectively in *L. cosentinii*, and 746,976, 428,940 and 101,168, respectively in *L. digitatus* (**Figure 4a,b**). The low scores in the tetraploid model therefore favor tetraploidy. The same analysis was applied to individual sequences from both species to determine whether some sequences or chromosomes have a different ploidy to the rest of the genome (a sign of aneuploidy). However, the lowest scores for all sequences again favored the tetraploid model (**Figure 4c,d**). Our data indicate that *L. cosentinii* and *L. digitatus* are tetraploid species with basic chromosome numbers *x* =8 and *x* =9, respectively, and confirm the absence of aneuploidy.

**Figure 3:**
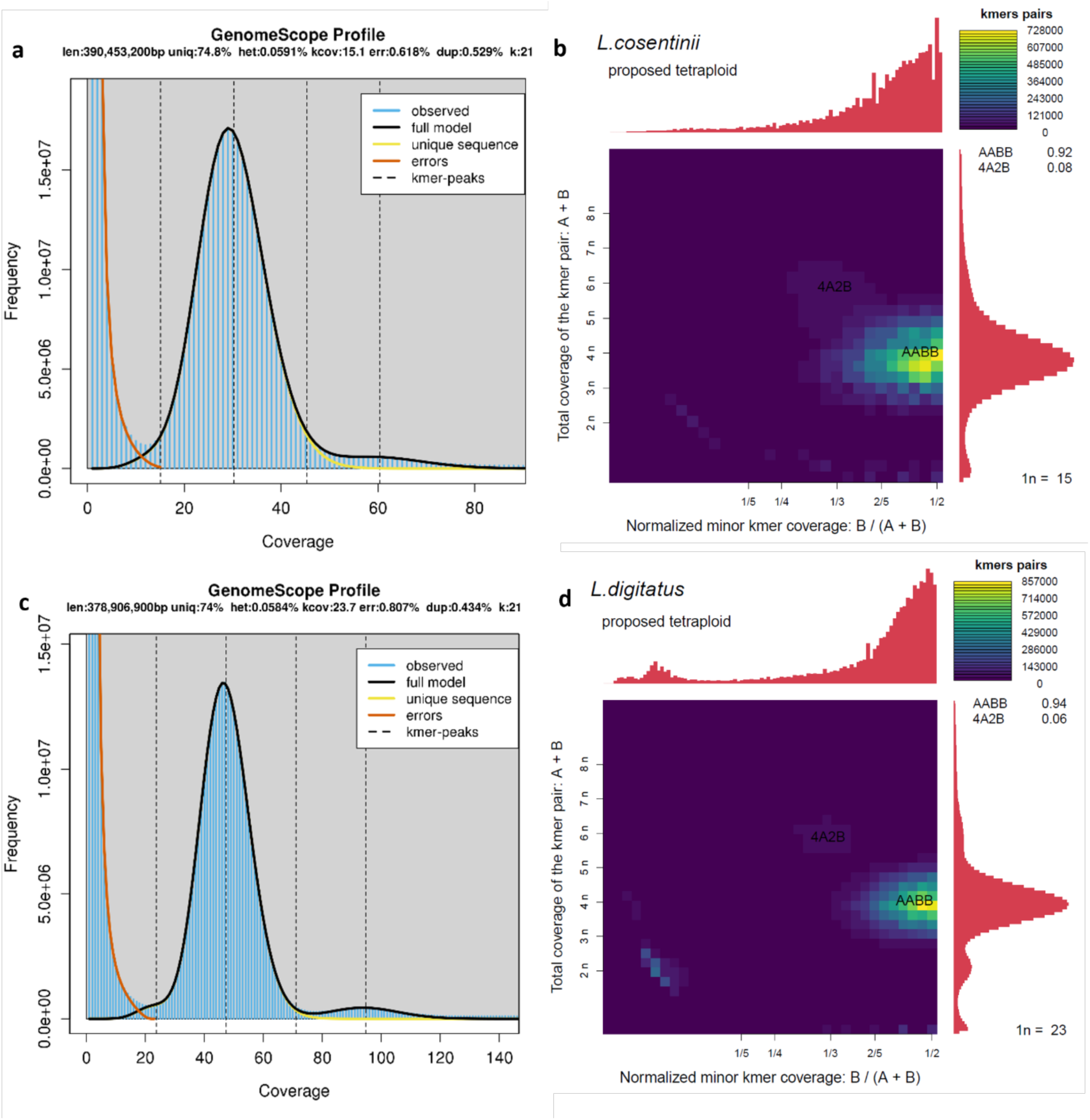
The k-mer distributions and proposed ploidy levels for *Lupinus cosentinii* and *L. digitatus*. (a) *L. cosentinii* GenomeScope profile. (b) *L. cosentinii k*-mer distribution. (c) *L. digitatus* GenomeScope profile. (d) *L. digitatus k*-mer distribution.

**Figure 4:**
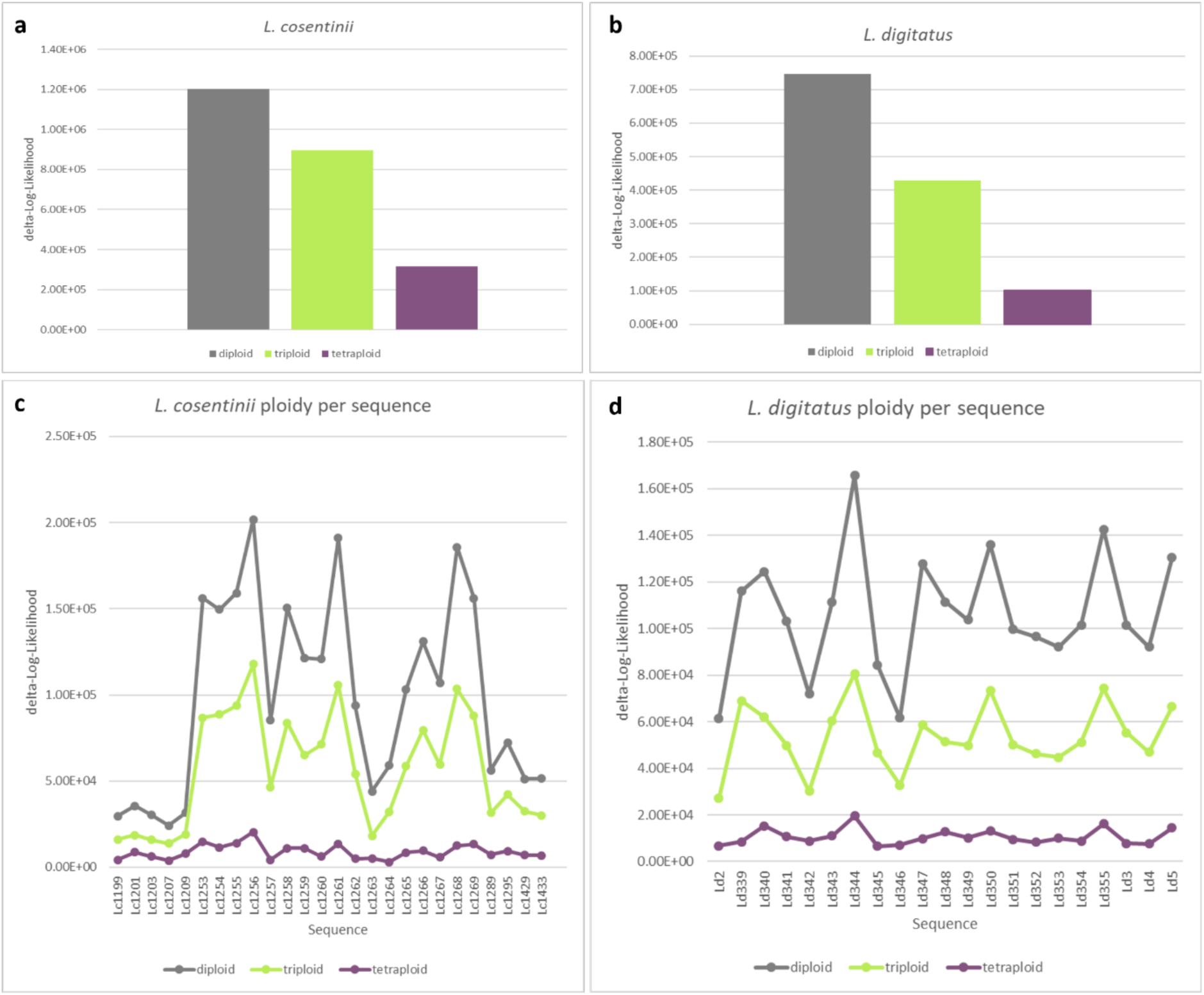
Ploidy prediction in *Lupinus cosentinii* and *L. digitatus* according to nQuire. The lower the delta log-likelihood score, the better the fit to the corresponding model. (a) Prediction scores on the *L. cosentinii* whole genome assembly and (c) on the longest 26 sequences (N90). (b) Prediction score on the *L. digitatus* whole genome assembly and (d) on the longest 21 sequences (N80).

Today’s legumes are proposed to have originated from a legume-common tetraploid (LCT) ancestor that existed ∼60 Ma (Schmutz et al. 2010, Wang et al. 2017), suggesting that tetraploid crops such as peanut and lupin may have originated from the same whole-genome duplication (WGD) event shared with most papilionid species (Bertioli et al. 2009, Doyle 2012). Alternatively, WGD may have occurred in a *Lupinus* diploid ancestor after the lupins diverged from the remaining Genisteae 19–35 Ma. Polyploidy has also been reported in two lupin crops (*L. angustifolius* and *L. albus*) but the basic chromosome number remains ambiguous. In *L. albus*, hexaploidization has been proposed based on a WGD event ∼60 Ma followed by a whole-genome triplication (WGT) event 22–25 Ma (Hane et al. 2017, Xu et al. 2020), and. This WGT, shared by *L. albus* and *L. angustifolius* as ‘extra’ duplications, may be linked to a higher frequency of polyploidization in domesticated species compared to wild relatives (Salman-Minkov et al. 2016).

The WGD/WGT events in the early genistoid clade are questionable and only partially supported (Cannon et al. 2015). The complex evolution of the *Lupinus* genus is characterized by remarkable diversity in genome size, basic and somatic chromosome numbers, and chromosome rearrangements, in contrast to other legume genera. For example, *Phaseolus* and *Cajanus* (phaseoloids) feature mostly diploid species with the same chromosome number 2*n* = 22, whereas *Arachis* (dalbergioids) features both diploids (2*n* = 20) and tetraploids (2*n* = 40), and *Dalbergia* (dalbergioids) features exclusively diploid species with the chromosome number 2*n* = 20 (Hung et al. 2020). Interestingly, the *Lupinus* diploid ancestor with a basic chromosome number of *x* = 9 (Xu et al. 2020) was confirmed across the genistoids (Doyle 2012). However, the basic chromosome number may differ in early-diverging genistoid species, including those in the genus *Sophora* such as *S. flavescens* (2*n* = 18, diploid) (Qu et al. 2023) and *S. japonica* (2*n* = 28, ploidy unreported) (Lei et al. 2022), and also *Crotalaria* spp. (2*n* = 14, 16 or 32) (Mondin and Aguiar-Perecin 2011). Furthermore, the genistoid genus *Ulex* has a chromosome number of 2*n* = 32, 64 or 96 and *Genista* has a chromosome number of 2*n* = 48, 44 (described as aneuploid), 72 or 96 (Bacchetta et al. 2012). *L. digitatus* (2*n* = 36) is the only known Old World lupin that provides a direct example of *x* = 9, corresponding to the *Lupinus* diploid ancestor. In contrast, *L. cosentinii* has a different basic chromosome number (*x* = 8) and refutes the hypothesis that species with chromosome numbers such as 2*n* = 32 are aneuploids (Nevado et al. 2016, Susek et al. 2019). However, *x* = 8 is considered a primitive basic number of the genistoids.

### Comparative genomics in lupin species

We compared our *L. cosentinii* and *L. digitatus* genome assemblies to the published *L. albus* genome (Hufnagel et al. 2020), which is the most annotated and curated genome assembly in the *Lupinus* genus. Systematic pairwise comparisons revealed large syntenic blocks conserved in all three genomes (**Figure 5**). The largest blocks were 24.3, 19.9 and 17.3 Mb in length, consisting of 1611, 747 and 1130 collinear genes in the *L. digitatus* vs *L. cosentinii* (LdLc), *L. digitatus* vs *L. albus* (LdLa) and *L. albus* vs *L. cosentinii* (LaLc) comparisons, respectively (**Figure 6**). The degree of duplication showed a similar distribution when considering the total number of genes and genes located in smaller syntenic blocks. The average duplication level of the total number of genes was similar in *L. cosentinii* (1.32) and *L. digitatus* (1.28) but increased to 1.43 in both species when considering the four smaller syntenic blocks (**Table 3**). The K_a_/K_s_ ratios in all three comparisons were < 1, showing that most genes are under stabilizing selection. LaLc and LdLa showed similar distributions, in both cases narrower than that of LdLc (**Figure 7**). The differences were highlighted when looking at the K_s_ distributions, where LaLc and LdLa showed similar K_s_ values, both significantly higher than LdLc.

**Figure 5:**
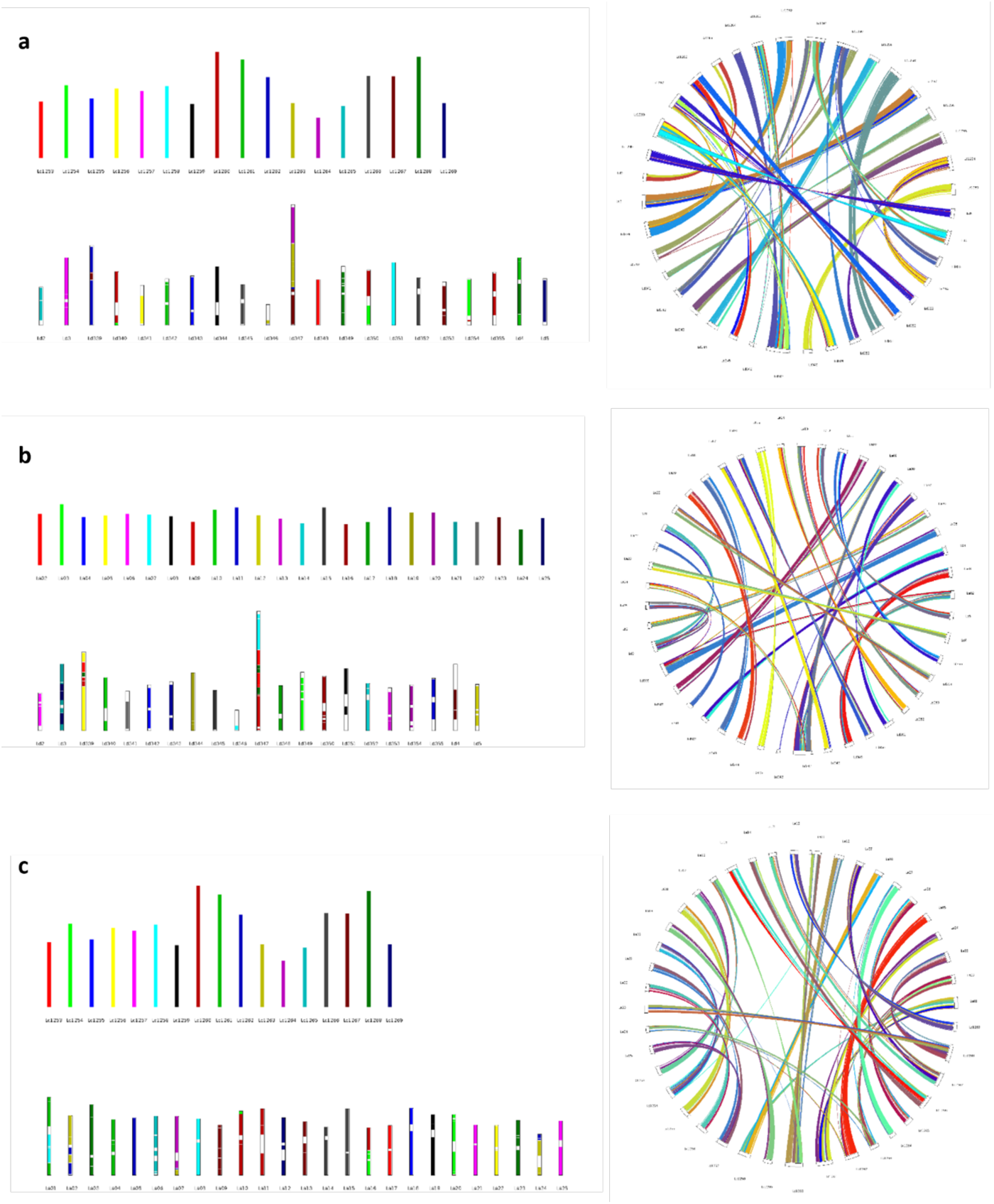
Collinearity analysis. (a) Collinear blocks between the assembled genomes of *Lupinus digitatus* (Ld) and *L. cosentinii* (Lc). (b) Collinear blocks between the assembled genomes of *L. albus* (La) and *L. digitatus* (Ld). (c) Collinear blocks between the assembled genomes of *L. albus* (La) and *L. cosentinii* (Lc).

**Figure 6:**
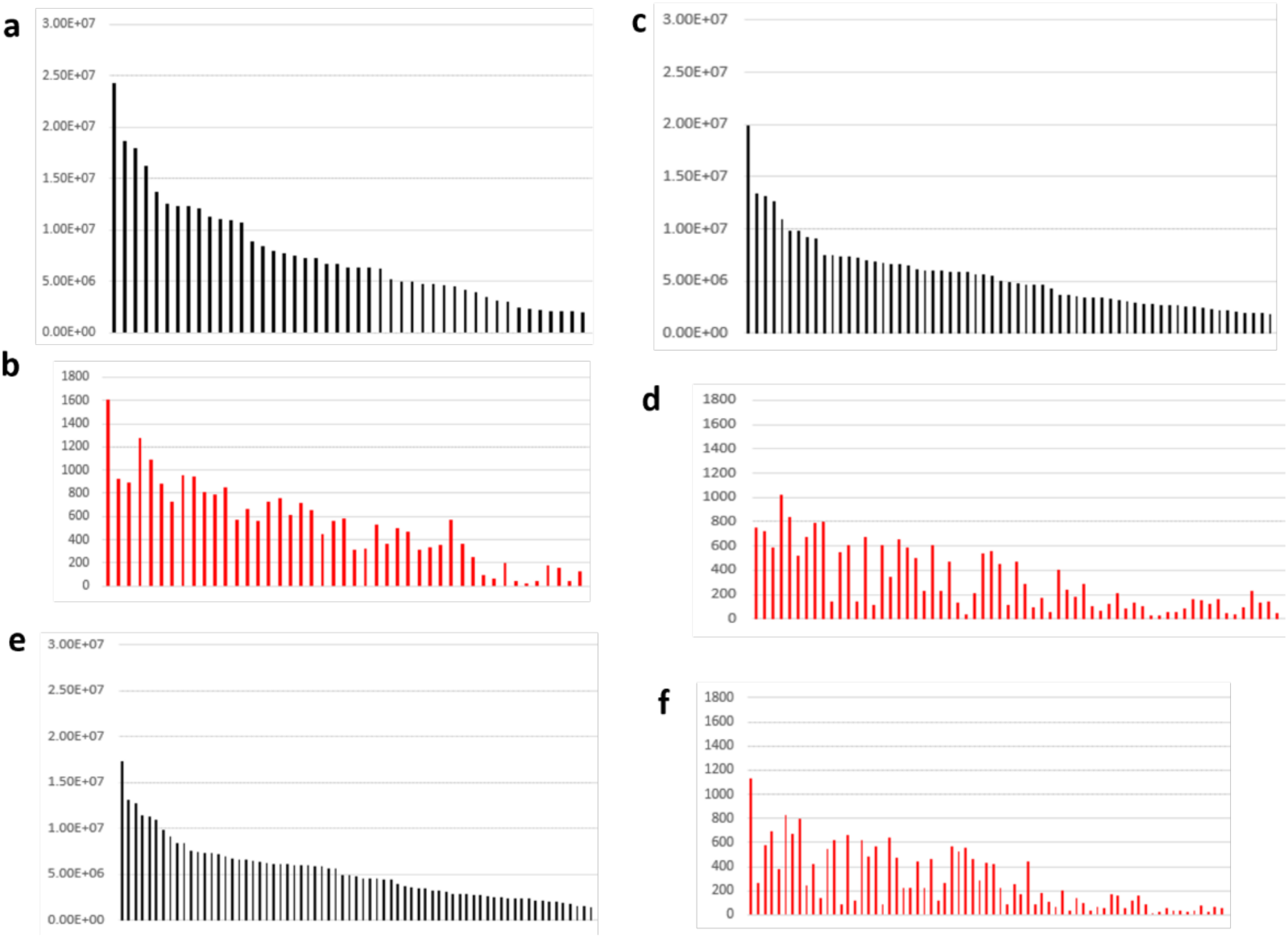
Size in Mb of syntenic blocks and number of collinear genes in syntenic blocks between lupin genomes. (a) *Lupinus digitatus* and *L. cosentinii* syntenic blocks and (b) collinear genes. (c) *L. digitatus* and *L. albus* syntenic blocks and (d) collinear genes. (e) *L. albus* and *L. cosentinii* syntenic blocks and (f) collinear genes.

**Figure 7:**
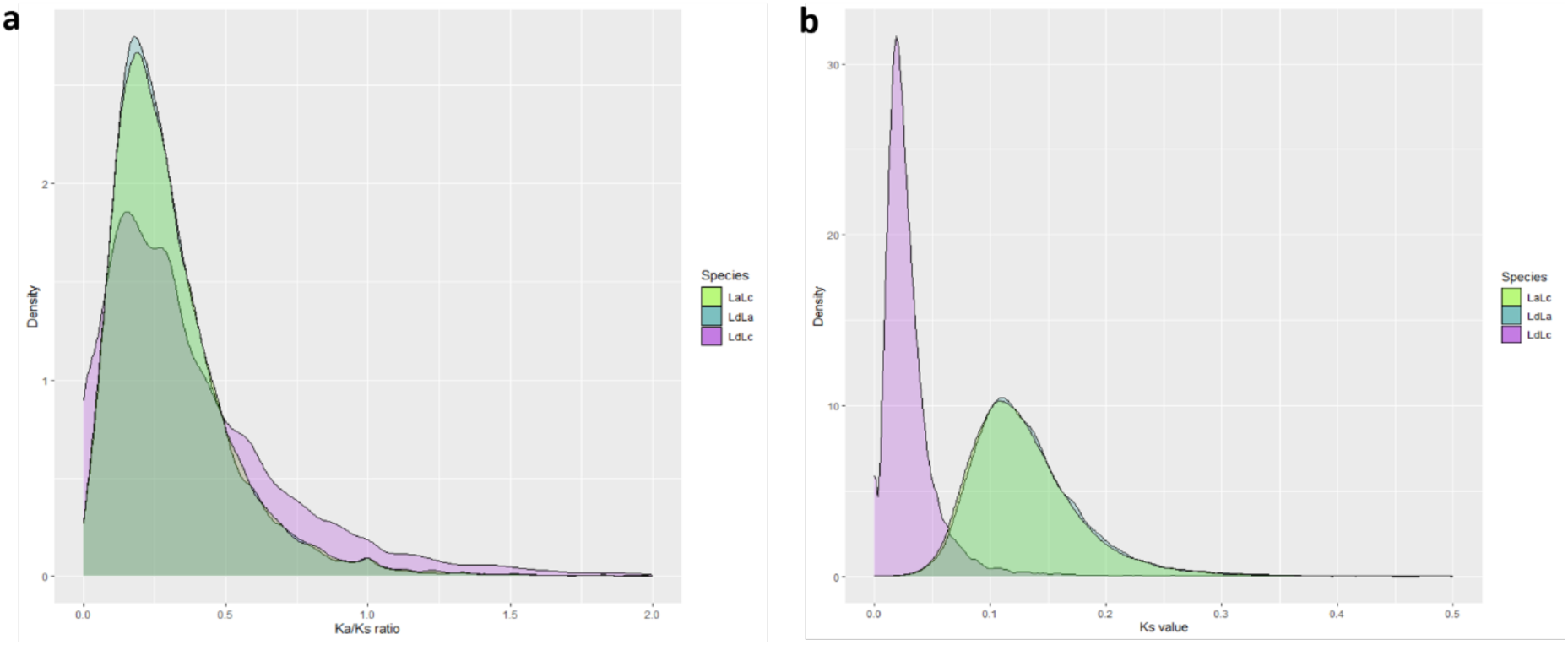
Distribution of (a) K_a_/K_s_ ratio and (b) K_s_ score calculated from the coding sequences of orthologous genes from the three comparisons: *Lupinus. digitatus* and *L. cosentinii* (LdLc); *L. digitatus* and *L. albus* (LdLa); *L. albus* and *L. cosentinii* (LaLc).

**Table 3:** Average degree of gene duplication in *Lupinus cosentinii* and *L. digitatus* when considering the total number of genes and when focusing on genes in small syntenic blocks.

| Average duplication level: | *L. cosentinii* | *L. digitatus* |
| --- | --- | --- |
| All genes | 1.32 | 1.28 |
| Genes in small syntenic blocks | 1.43 | 1.43 |

We clustered genes into 25,663 families in *L. cosentinii*, *L. digitatus* and *L. albus* (reference species). Most of the gene families (21,273) were shared by all three species (**Figure 2c**), but 778 were shared only between *L. cosentinii* and *L. albus* and 561 only between *L. digitatus* and *L. albus*. The number of single-species gene families was similar in *L. cosentinii* and *L. digitatus* (94 and 96, respectively). The numbers of expanded gene families (1035 and 1089, respectively) and contracted gene families (3518 and 3663, respectively) was also similar in these species (**Table 4**). GO enrichment showed that the expanded gene families in *L. cosentinii* were enriched in 39 terms such as DNA integration, lipid transport and phloem development, whereas those in *L. digitatus* were enriched in 57 terms such as defense responses and sucrose transportation (**Supplementary Table S6**).

**Table 4:** Summary of expanded and contracted gene families, number of genes in families, and genes lost from *Lupinus albus* a reference species.

|  | Expanded |  |  | Contracted |  |  |
| --- | --- | --- | --- | --- | --- | --- |
|  | Gene families | Genes | Genes gained relative to *L. albus* | Gene families | Genes | Genes lost from *L. albus* |
| *L. cosentinii* | 1305 | 3076 | 1711 | 3518 | 2644 | 4978 |
| *L. digitatus* | 1089 | 2633 | 1497 | 3663 | 2629 | 5182 |

The expanded gene families from the two species featured three common enriched GO terms related to DNA organization: double-stranded DNA binding, enzyme inhibitor activity, and nucleosome. These families were investigated in more detail, seeking evidence for the expansion or contraction of genes involved in seed size in *L. albus*. albus (**Supplementary Table S7**). We identified 12 gene families whose gene number had doubled in *L. cosentinii* and *L. digitatus* compared to *L. albus*. However, we also found some genes that were only present in *L. cosentinii*, including gene families involved in cell division (six genes in *L. cosentinii* compared to only three in *L. albus*) and in seed development/growth (two genes in *L. cosentinii* compared to only one in *L. albus*). The latter genes therefore appear to play important roles in *L. cosentinii* and *L. albus* but not in *L. digitatus*, probably in the context of environmental stress responses. The common expanded gene families in *L. cosentinii* and *L. digitatus* (double the gene number compared to *L. albus*) were related to sugar nucleotide biosynthesis (UDP-D-glucuronate 4-epimerase), sugar metabolism and embryo cellularization. These are likely to regulate seed size by controlling the size of the embryo, and may affect seed development in response to environmental stress (e.g., high temperature). The timing of embryo cellularization is determined by the phosphatidylethanolamine-binding protein (PEBP) family, which comprises the FLOWERING LOCUS T (FT)-like, TERMINAL FLOWER1 (TFL1)-like, and MOTHER OF FT AND TFL1 (MFT)-like subfamilies (Karlgren et al. 2011), and the number of PEBP-like genes in plants varies greatly, (Wang et al. 2015). The UDP-glucuronate 4-epimerases gene families were also expanded in *L. cosentinii* and *L. digitatus*, and are also expanded in the legume cluster ‘bean’ (Gaikwad et al. 2023). In the analysis of over-represented GO terms for expanded gene families, we found enrichment for ‘molecular function’ GO categories related to signaling receptor activity, abscisic acid (ABA) binding, protein phosphatase inhibitor activity, triglyceride lipase activity, and glucose-1-phosphate adenylyltransferase activity in *L. cosentinii* and *L. digitatus* but not *L. albus*. The adaption of these species to drought stress may therefore involve more efficient ABA signaling (Zhu 2016). Triglyceride lipase and protein phosphatase inhibitor activity may also improve drought tolerance in plants (Xiang et al. 2017). These expanded gene families in *L. cosentinii* and *L. digitatus* may be the remnants of a lineage-specific second duplication contemporaneous with species divergence. Triglyceride lipase is associated with lipid degradation (Rosnitschek and Theimer 1980), but a comparative lipidomic analysis of *L. cosentinii*, *L. digitatus* and *L. albus* is needed to explore the relevance of this observation. Glucose-1-phosphate adenylyltransferase activity is important in starch and sucrose metabolism, and the expanded gene families (along with UDP-D-glucuronate 4-epimerase) may contain novel genes that determine seed size and quality specifically in *L. cosentinii* and *L. digitatus*.

We speculate that these sets of genes play a fundamental role in lupin seed size determination and have remained intact after genome duplication. Therefore, our genomic data will help to decipher the evolutionary pathways and underlying mechanisms, which will facilitate the development of new breeding strategies.

## Conclusions

We have described the first whole-genome assemblies of the wild lupin species *L. cosentinii* and *L. digitatus*, providing insight into lupin genomics and evolution, and adding to the genetic resources available for lupin breeding and crop improvement. These two annotated assemblies provide key reference genomes for lupin and the genistoid clade more generally. Importantly, we provide evidence that both species are tetraploid. Our data provide insight into the role of genome duplication in lupin evolution but further evidence from other wild and domesticated species would help to complete the picture. *L. cosentinii* and *L. digitatus* are the only wild lupin species whose ploidy has been confirmed, and this will help us to understand the domestication, agricultural improvement, environmental adaptability and evolution of crops, facilitating the exploitation of legume plants as part of a healthy and sustainable diet (Bellucci et al. 2021, Nadon and Jackson 2020). As well as providing a source of dietary protein, lupins also have many advantages in the context of legume-based cropping systems, including up to 40% higher yields in subsequent rotations (Zhao et al. 2022).

Interestingly, one of the main groups of New World lupins has a chromosome number of 2*n* = 36, although exceptionally *L. bracteolaris*, *L. linearis* and others have a chromosome number of 2*n* = 32, like *L. cosentinii* (Maciel and Schifino-Wittmann 2002). This coincidental matching chromosome number in Old and New World lupins may raise questions about the geographical origin of lupins, which is proposed to be in the Old World (Nevado et al. 2016). In the northern hemisphere, where the genistoid clade is thought to have originated during the Paleocene epoch (Doyle 2012), evolutionary studies of these two groups of lupins will shed new light on the evolution of the entire *Lupinus* genus.

The analysis of lupin gene families provided insights into their relationship with phenotypic diversification and species adaptation, which will facilitate the exploitation of underutilized legume species by identifying genes that can be used in crop breeding programs. The expansion and contraction of gene families involved in seed size, a paradigmatic domestication trait, indicate that gene duplication may have led to morphological adaptations in *L. cosentinii* and *L. digitatus* differing from those in *L. albus*, although additional work is needed to validate these hypotheses. Seed size may therefore reflect convergent evolution mechanisms that play a key role in crop domestication.

## Methods

### Plant materials

The characteristics of *L. cosentinii* and *L. digitatus* are summarized in **Table 5 and shown in Figure 2d** . The source material was used to develop single-seed descent lines (at least two rounds of multiplication). Seeds of both species were scarified, vernalized for 21 days and then sown in 7.5-L pots containing a 1:1 mix of peat and vermiculite. The plants were grown in a phytotron at 22/18 °C (day/night temperature) with a 16-h photoperiod, 60–65% relative humidity, and watering as required.

**Table 5.** Key characteristics of *Lupinus cosentinii* and *L. digitatus*.

|  | *L. cosentinii* | *L. digitatus* |
| --- | --- | --- |
| ID, provider | 98640, Polish *Lupinus* Gene Bank, Breeding Station Wiatrowo, Poznan Plant Breeders Ltd, Poland | PI 660697, US Department of Agriculture, USA |
| Genome size (Mb), flow cytometry | 695.80 | 671.30 |
| Chromosome number ($2n$) | 32 | 36 |
| Geographic distribution | western Mediterranean coast | pan-Saharan region |

### Extraction of high-molecular-weight DNA

For PacBio sequencing, high-molecular-weight DNA was extracted from 1 g frozen young leaf material that had been ground to powder under liquid nitrogen. Nuclei were isolated in NIBTM buffer (10 mM Tris, 10 mM EDTA, 0.5 M sucrose, 80 mM KCl, 8% PVP (MW 10 kDa), 100 mM spermine, 100 mM spermidine, pH 9.0) supplemented with 0.5% Triton X-100 and 0.2% 2-mercaptoethanol followed by filtration through 100-µm and 40-µm cell strainers and centrifugation (2500 g, 10 min, 4 °C) as previously described (Zhang et al. 2012). DNA was then extracted from nuclei using the Genomic-tip 100/G kit (Qiagen) and eluted in low TE buffer (10 mM Tris, 0.1 mM EDTA, pH 9.0). DNA size and integrity were analyzed by pulsed-field gel electrophoresis (PFGE) using the CHEF Mapper system (Bio-Rad Laboratories) with a 5–450 kb run program. DNA was quantified using the Qubit DNA BR Assay Kit and a Qubit fluorimeter (Thermo Fisher Scientific) and its purity was evaluated by spectrophotometry using a Nanodrop 2000 (Thermo Fisher Scientific). PacBio libraries were prepared from both species using the SMRTbell prep kit v3.0, followed by SMRT sequencing on a Sequel II device, generating ∼32.8 and 18.9 Gb for *L. cosentinii* and *L. digitatus*, respectively

### Whole-genome library preparation for Illumina sequencing

We fragmented 700 ng of high-molecular-weight DNA using a Covaris S220 sonicator and a WGS library was generated for both species using the KAPA Hyper Prep kit with a PCR-free protocol according to the manufacturer’s instructions (Roche). We applied final size selection using a 0.7-fold ratio of AMPureXP beads. The sequence length was assessed by capillary electrophoresis on a Tape Station 4150 (Agilent Technologies) and the library was quantified by qPCR against a standard curve with the KAPA Library Quantification Kit (Kapa Biosystems). Libraries were sequenced on a NovaSeq6000 Illumina platform in 150PE mode, generating ∼130 and ∼124 million fragments for *L. cosentinii* and *L. digitatus*, respectively.

### Extraction of ultra-high-molecular-weight DNA and Bionano optical mapping

DNA was extracted from fresh *L. cosentinii* and *L. digitatus* sprouts/leaves < 2 cm in length, which had been kept in the dark for ∼16 h before extraction (Canaguier et al. 2022). DNA was prepared from ∼0.4 g of tissue using the Bionano Prep High Polysaccharides Plant Tissue DNA Isolation Protocol (document number 30128 revision C) with 2-mercaptoethanol in the wash buffer. Two plugs were prepared for each species. DNA extracted from one plug was used for quality assessment by PFGE with the CHEF Mapper system (5–450 kb run program, two-state mode, 20 h). DNA was quantified using the Qubit DNA BR Assay Kit and a Qubit fluorimeter as above. DNA from the other plug was used for Bionano optical mapping on a Bionano Saphyr, generating 4.7 and 5.4 million molecules for *L. cosentinii* and *L. digitatus*, respectively.

### Hi-C library preparation for Illumina sequencing

Libraries were prepared from 0.52 frozen young *L. cosentinii* and *L. digitatus* leaves using the Proximo Hi-C (Plant) Kit and protocol v4.0 (Phase Genomics) with three extra wash steps and 12 PCR amplification cycles. Hi-C libraries were analyzed by capillary electrophoresis on a Tape Station 4150 using a D1000 ScreenTape (average size 469 bp, concentration 9.40 ng/µl in 30 µl). Libraries were quantified by real-time PCR against a standard curve with the KAPA Library Quantification Kit and were sequenced on a NovaSeq 6000 Illumina platform in 150PE mode, generating 201 and 354 million fragments for *L. cosentinii* and *L. digitatus*, respectively.

### RNA-Seq library preparation for Illumina sequencing

We prepared 22 RNA samples from *L. cosentinii* (six samples of small leaves, four of big leaves, four leaf stem, four pod and four root) as well as 32 samples from *L. digitatus* (six of leaves, six leaf stem, six pod, five stem, four apical stem, three lateral root and two main root). Total RNA was isolated from 30 mg of ground plant tissue using the SV Total RNA Isolation System Kit (Promega) and its concentration and integrity were assessed using the RNA 6000 Nano Kit on a Bioanalyzer (Agilent Technologies). All samples showed an RNA integrity number (RIN) > 7. Samples were quantified using the Qubit RNA HS Assay Kit (Thermo Fisher Scientific). We pooled 2–3 RNA samples from the same tissue for library preparation to make pools of five different *L. cosentinii* tissues (small leaf, big leaf, leaf stem, pod and root) and seven different *L. digitatus* tissues (leaf, pod, stem, lateral root, main root, leaf stem and apical stem). RNA-Seq libraries were generated using the TruSeq stranded mRNA ligation kit (Illumina) from 1000-ng RNA samples, after poly(A) capture and according to the manufacturer’s instructions. Library quality and size were assessed by capillary electrophoresis using the Tape Station 4150. Libraries were quantified by real-time PCR against a standard curve with the KAPA Library Quantification Kit. The libraries were pooled at equimolar concentrations and sequenced on a Novaseq6000 device in 150PE mode yielding ∼30 million fragments per sample.

### *De novo* genome assembly from PacBio Hi-Fi reads

PacBio Hi-Fi reads were assembled *de novo* using HiCanu v2.1.1 (Nurk et al. 2020) with default parameters. Completeness was evaluated using BUSCO v5.4.7 (Manni et al. 2021) and the Fabales_obd10 database comprising of 5366 genes. Illumina WGS data were evaluated using FastqQC v0.11.9 and low-quality segments and sequencing adapters were removed using Fastp v0.21.0 (Chen et al. 2018). Filtered reads were aligned on the assembly using bwa-mem2 v0.7.17 and residual base-level errors were corrected by three rounds of polishing using Pilon v1.23 . To evaluate the effectiveness of this approach, we applied variant calling using GATK HaplotypeCaller v4.2.2 (McKenna et al. 2010) before and after polishing. We also used purge_haplotigs v1.1.2 (Roach et al. 2018) to remove putative haplotype duplications. BLAST v2.9.0+ (Altschul et al. 1990) was used to screen all remaining reads against the NCBI nt database to confirm that all reads belonged to the kingdom Viridiplantae, thus ensuring the absence of contamination. BLAST was also used to screen mitochondrial (Link1) and chloroplast (Link2) RefSeq databases and published *L. albus* organelle sequences (Hufnagel et al. 2020) to exclude organelle DNA.

### Scaffolding with Bionano optical maps

Bionano sequencing outputs were filtered to remove molecules < 150 kb in length before *de novo* assembly and alignment on the corresponding genome maps using Bionano Solve v3.7.1 (Link3).

### Chromosome-level scaffolding with Hi-C data

The Hi-C raw reads were aligned on the Bionano genome assemblies using the Juicer v1.6 pipeline (Durand et al. 2016) before a second round of scaffolding using 3d-dna v18.09.22 (Dudchenko et al. 2017) with default parameters. Before the misjoin correction step, the Hi-C contact matrix was manually curated with juicebox v1.11.08.

### Structural annotation

A library of repeats was constructed using RepeatModeler v2.0.2 (Flynn et al. 2020) and repetitive sequences were soft masked using RepeatMasker v4.1.2-p (Tarailo-Graovac and Chen 2009). RNA-Seq data were aligned on the assembled genome using Hisat2 v2.2.1 (Kim et al. 2019) with a maximum intron length of 60 kb. The alignments were then converted into intronic hints, retaining only those supported by at least 10 reads. RNA-Seq data were also assembled into transcripts using Trinity v2.15 (Grabherr et al. 2011). Only the primary isoform of all reconstructed genes, namely those classified as ‘main’ and ‘complete’ by Evidential Gene v2018, were retrieved and aligned on the assembled genome using gmap v2017-11-15 (Wu and Watanabe 2005) for use as extrinsic evidence. Finally, proteins from the closely related species *L. albus* (Link4) were aligned on the genome assembly using Genome Threader v1.7.1 (Gremme et al. 2005). The extrinsic evidence extracted from the three different sources described above was then used for final *ab initio* gene prediction with Augustus v3.3.3 (Hoff and Stanke 2019) trained using *Fabales* BUSCO genes (BUSCO v5.4.7, Fabales_odb10 database). The predicted genes were filtered using Interproscan v5.52-86.0 (Jones et al. 2014) to retain only genes with known protein domains.

### Functional annotation

Genes were functionally annotated based on the analysis of homology (BLAST v2.9.0+, keeping only the best hits for each gene) and protein domains (Interproscan). For homology-based analysis, we considered three levels of confidence: (1) genes with functional annotations in SwissProt (Link5), RefSeq plant databases (Link6) and/or TAIR (Link7) were labeled as high confidence; (2) genes were labeled as medium confidence if we retrieved functional annotations based on the *L. albus* proteome; and (3) genes that were not annotated using the first two levels were screened against the NBCI nr database to obtain a descriptive annotation. The alignments were filtered by percentage coverage and identity, both with thresholds of 80%. GO terms were derived from homology-based analysis at the first and second confidence levels (if the function was concordant) and from Interproscan analysis.

### Ploidy analysis

The level of ploidy in *L. cosentinii* and *L. digitatus* was assessed using two methods, the first based on *k*-mer distribution and the second on biallelic SNP frequencies, applied to Illumina reads after noise reduction. For the first approach, the *k*-mers in WGS Illumina reads were counted using KMC v3.2.2 (Kokot et al. 2017). The *k*-mer distributions were analyzed using Genomescope2.0 and Smudgeplot (Ranallo-Benavidez et al. 2020) with parameter –homozygous due to the high level of homozygosity. For the second approach, Gaussian mixture models were used to estimate the ploidy level with nQuire (Weiß et al. 2018). Sequencing reads were mapped to the genome and biallelic SNP frequencies were calculated. A delta log-likelihood score was then calculated between a free model and three fixed models (diploid, triploid and tetraploid). The lowest fixed-model score points to the most likely ploidy. This analysis was applied to the whole *L. cosentinii* and *L. digitatus* datasets and also to 26 individual sequences in *L. cosentinii* (corresponding to ∼80% of the genome assembly) and 21 in *L. digitatus* (corresponding to ∼90% of the genome assembly). If some sequences showed a lower score in a different ploidy model than the rest of the genome, this could be interpreted as a sign of aneuploidy. The rationale behind the use of two approaches was to validate the predicted ploidy level independently from the homozygosity of the two assembled genomes.

### Comparative genomics

Synteny was evaluated using MCScanX with default parameters. Specifically we used MCScanX_h, allowing the exploitation of orthologous genes from *L. albus*, *L. cosentinii* and *L. digitatus* predicted by Orthofinder v2.5.4 (Emms and Kelly 2019). We tested the pairwise comparisons *L. digitatus* vs *L. cosentinii* (LdLc), *L. digitatus* vs *L. albus* (LdLa) and *L. cosentinii* vs *L. albus* (LcLa). To assess the relationship between any two species, we analyzed the degree of duplication considering the total number of genes or only genes located in four smaller blocks of synteny (Lc1254&Ld340, Lc1267&Ld339, Lc1263&Ld346 and Lc1260&Ld354). We analyzed 304 genes each for *L. cosentinii* and *L. digitatus*. Genes in an orthogroup from the same species were considered as paralogs and thus an indication of gene duplication. The paralogous genes in *L. cosentinii* and *L. digitatus* were predicted using Orthofinder v2.5.4. The coding regions of the orthologous gene pairs from the three pairwise comparisons were used to calculate *K_a_/K_s_* ratios in the MCScanX downstream analysis package add_kaks_to_synteny. Variation in gene family sizes was analyzed using Cafe5 (Mendes et al. 2020) followed by GO functional enrichment analysis of the expanded gene families.

## Data availability

The raw sequence reads generated and analysed in this study have been deposited in the Sequence Read Archive (SRA) of the National Center of Biotechnology Information (NCBI) under BioProject IDs: PRJNA1080360 (*L. cosentinii*) and PRJNA1080354 (*L. digitatus*); and Biosample IDs : SAMN40127157 (*L. cosentinii*) and SAMN40126867 (*L. digitatus*).

## Author contributions

K.S. conceptualized the study, designed the experiments, wrote the manuscript and supervised the project. E.F. conducted the bioinformatic analysis and helped to write the manuscript. M.T. cultivated the plants under controlled conditions, M.K. analyzed the plants, extracted nucleic acids, and assisted with data interpretation. H.J. and U.T. assisted with bioinformatic analysis and manuscript preparation. M.N.N., R.P. and S.A.J. assisted with data interpretation. R.P., M.D. and S.A.J. and contributed to the editing and substantive revision of the manuscript. All authors read and approved the final version of the manuscript.

## Links

**Link1:** https://ftp.ncbi.nlm.nih.gov/refseq/release/mitochondrion/

**Link2:** https://ftp.ncbi.nlm.nih.gov/refseq/release/plastid/

**Link3:** (https://bionanogenomics.com/support/)

**Link4:** (https://phytozome-next.jgi.doe.gov/info/Lalbus_v1)

**Link5:** https://www.uniprot.org/uniprotkb?facets=reviewed%3Atrue&query=%2A

**Link6**: https://ftp.ncbi.nlm.nih.gov/refseq/release/plant/

**Link7:** https://www.arabidopsis.org/download_files/Proteins/TAIR10_protein_lists/TAIR10_pep_20101 214

